# Initial pig developmental stage influences intestinal organoid growth but not phenotype

**DOI:** 10.1101/2024.01.26.577507

**Authors:** Camille Duchesne, Gwénaëlle Randuineau, Laurence Le Normand, Véronique Romé, Samia Laraqui, Alexis Pierre Arnaud, Gaëlle Boudry

## Abstract

Intestinal organoids are promising tools in the context of animal experiment reduction. Yet, a thorough characterization of the impact of the origin of intestinal stem cells (ISC) on organoid phenotype is needed to routinely use this cellular model. Our objective was to evaluate the effect of ISC donor age on the growth, morphology and cellular composition of intestinal organoids derived from pig, a valuable model of Humans. Organoids were derived from jejunal and colonic ISC obtained from 1, 7, 28, 36 and 180-day old pigs and passaged three times. We first confirmed by qPCR that the expression of 18% of the >80 studied genes related to various intestinal functions differed between jejunal and colonic organoids after two passages (P<0.05). Growth and morphology of organoids depended on intestinal location (greater number and larger organoids derived from colonic than jejunal ISC, P<0.05) but also pig age. Indeed, when ISC were derived from young piglets, the ratio of organoids to spheroids was greater (P<0.05), spheroids were larger during the primary culture but smaller after two passages (P<0.05) and organoids smaller after one passage (P>0.05) compared to ISC from older pigs. Finally, no difference in cellular composition, evaluated by immunostaining of markers of the major intestinal cell types (absorptive, enteroendocrine and goblet cells) were observed between organoids originating from 7 or 180-day old pigs, while difference between intestinal site origin were noticed. In conclusion, while the age of the tissue donor affected organoid growth and morphology, it did not influence their phenotype.

## Introduction

The pig has been used for biomedical research for a long time due to its close physiology to that of Humans [1]. This is particularly true for the gastro-intestinal tract whose functions are close to that of Humans [2]. Yet, the use of preclinical models in research is nowadays largely questioned and the development of alternatives to animal models is necessary. Hence, intestinal organoids, self-renewing 3D-structures that recapitulate features of intestinal epithelium architecture and functions, are promising tools. Several groups established porcine intestinal organoid protocols (for example [3,4]). Yet, the role of the different factors influencing porcine intestinal organoid growth and phenotype still need to be addressed to routinely use them in research.

The intestinal epithelium is renewed every 3 to 5 days [5]. This perpetual movement of intestinal epithelial cells (IEC) is driven by the continuous asymmetric division of intestinal stem cells (ISC) nested at the bottom of intestinal crypts. These LGR5+ cells generate progenitor cells, characterized by the presence of Sox9 [6]. These cells then differentiate into different cell lineages as they migrate up the crypt-villus axis in the small intestine and up the crypt in the colon. A large proportion (80%) of cells differentiate into absorptive cells, characterized by specific transporters and enzymes such as sucrase-isomaltase (SI) in the small intestine and carbonic anhydrase (CA) in the colon, while the others differentiate into secretory cells, represented by goblet and entero-endocrine cells, characterized by the presence of mucin 2 and chromogranin A, respectively. The fourth cell type, Paneth cells, is formed by progenitor cells which migrate towards the base of the crypts [7]. These cells intercalated between ISCs have a key role in microbial defense but also stem cell maintenance [8]. Their exact function is still controversial in pigs, leading to the name Paneth-cell like in this species [7].

The pig intestinal epithelium phenotype is influenced by many factors. Among them, IEC location along the intestinal tract is critical. The small intestine and the colon differ by their epithelial morphology, but also largely by their specific functions. Several studies highlighted that these functional differences are imprinted *in vivo* in ISC since organoids originating from pig small or large intestine ISC retain the function of their tissue of origin [3,4]. Another determinant of the intestinal epithelium phenotype is the piglet development stage during the fetal period but also during the neonatal period, including weaning. The weaning period in piglet is experienced as a physiological stress and induces many intestinal epithelium adaptations [9,10]. These changes are thought to be largely driven by changes in the luminal environment occurring at weaning, including both gut microbiota composition and nutrients modifications [11]. Mussard et al revealed that porcine organoids derived from pre-(21-day old) and post-weaning (35-day old) piglets harbored very close phenotypes, suggesting that the weaning-induced changes in luminal environment do not imprint piglet ISC [12]. Beside weaning, the neonatal period (from birth to post-natal day 28 in pigs), is also a key period in terms of intestinal development, with changes in epithelium morphology and gene expression with post-natal age [13], as well as ISC dynamics depending on gut location [14]. As opposed to weaning where the luminal environment does not seem to impact ISC, we recently demonstrated that changes in the luminal environment composition during the first week of life can imprint ISC, since organoids derived from the colon of antibiotic-treated vs. vehicle-treated piglets retained some, but not all, features of the changes observed *in vivo* at the epithelial level [15].

Our aim was therefore to evaluate if the age of the pigs used to generate intestinal organoids impacts their growth and phenotype, extending from early to adult life. We first verified that organoids derived from jejunum and colon displayed distinct phenotype at the gene expression level. We then evaluated the growth and morphology of organoids generated from jejunum and colon of pigs at 5 different key developmental stages (1, 7, 28, 36 and 180 days of age) then focused on the phenotype of organoids derived from the jejunum and colon of 7 and 180-day old pigs at the protein level using an immunohistochemistry approach.

## Material and methods

### Experimental study design

Ethical approval was formally waived by the French Ministry of Higher Education and Research since this experiment involved only tissue collection after conventional slaughtering. Twenty Largewhite x Landrace x Pietrain pigs (INRAE UE3P, St-Gilles, France) were selected at key stages of the maturation process of intestinal epithelium: before weaning at 1, 7 and 28-day of age and after weaning at 36 and 180-day of age, with 4 animals per group (2 males, 2 females). Animals were anesthetized by electronarcosis and immediately followed by euthanasia by exsanguination. The design of the experiment is presented in Fig 1A. Pigs were raised as normal commercial animals. Noteworthy, piglets had access to solid feed from day 21 of age and were weaned at day 28.

**Fig 1:**
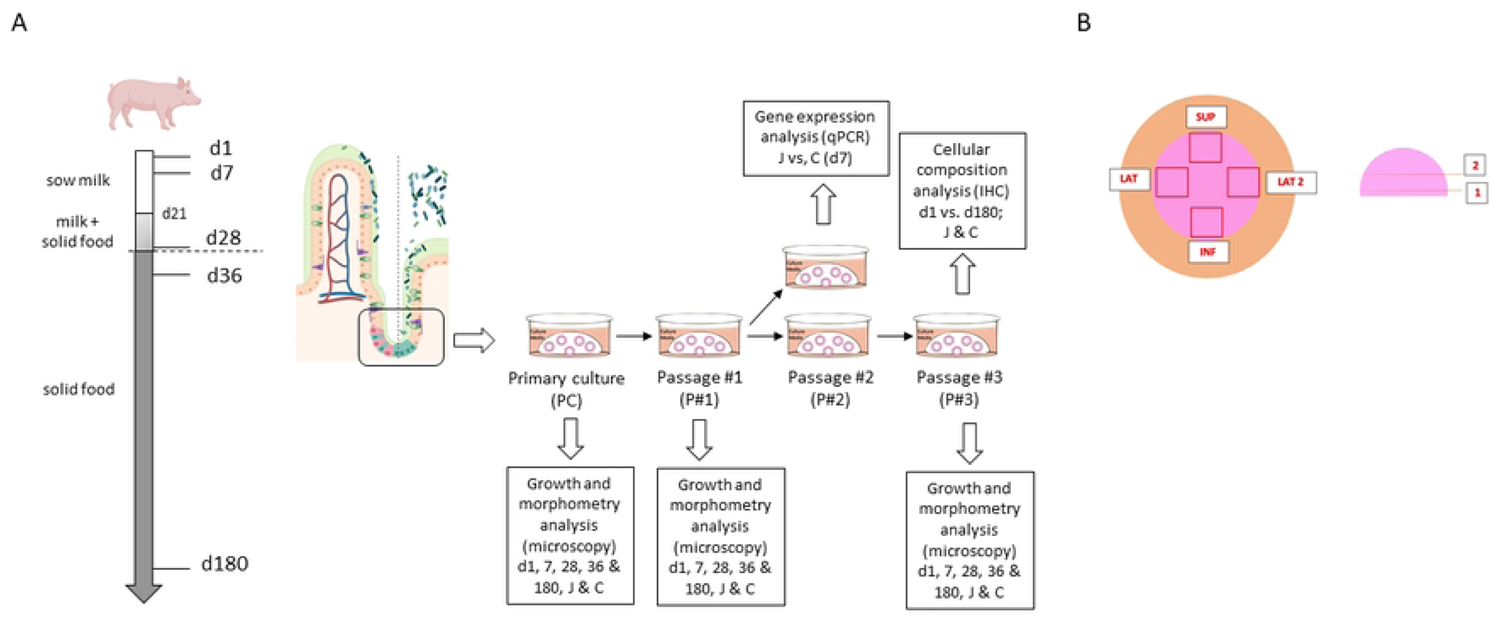
Overview of the experimental protocol. (A) Piglets were slaughtered at 1, 7, 28, 36 and 180 days of life to sample jejunal (J) and colonic (C) crypts that were used to produce organoids in a primary culture (PC) or after several passages (P#1, P#2, P#3) that were characterized using different techniques. (B) Schematic of the different pictures taken using inverted microscope to monitor organoid growth with time. Pictures of the superior, inferior and lateral quadrants were taken at two depth levels of the Matrigel dome.

### Intestinal tissue collection and preparation

For each pig, 5 cm sections of proximal colon and jejunum were isolated after euthanasia performed by electronarcosis then bleeding. Samples were thoroughly rinsed with phosphate buffer saline (PBS) supplemented with 5 mM of dithiothreitol (DTT, Sigma-Aldrich) and 1% of penicillin-streptomycin-amphotericine (PSA, Sigma-Aldrich) then transported within 10 min to the laboratory in a solution of PBS + 1% PSA on ice. The *muscularis* was removed for jejunum samples of pigs over 28 days of age. Each specimen was then opened longitudinally and pinned on a sylgard-coated petri dish, apical pole up. Villi were removed from the jejunum and mucus from the colon by scraping with angled forceps under a binocular magnifying glass. Tissues were then incubated for 30 min. at 80 rpm speed on a shaking plate in 10 mL of PBS supplemented with 3 mM DTT, 1% PSA, 9 mM EDTA (Sigma-Aldrich) and 10 μM Y27632 dihydrochloride (Rho-Kinase inhibitor, Preprotech). After incubation, the solution was removed and replaced by 10 mL of PBS supplemented with 1% PSA and 10 μM Y27632. Each sample was scraped using angled forceps under a binocular magnifying glass in order to release the crypts. The solution containing the crypts was immediately stored on ice.

### Three-dimensional organoid culture

The isolated crypts solutions were filtered at 200 μm then centrifuged at 100 g, 4° C for 5 min. Crypts were then suspended in 6 mL of cold Dulbecco’s Modified Eagle Medium (DMEM, Gibco) supplemented with 10 μM Y27632 and 1% PSA. Crypt density was determined in a 10 μL droplet with an optical microscope. 150 crypts were then seeded in 50 μL droplets of 4°C Matrigel Growth Factor Reduced Matrigel (Corning) in pre-warmed 24 well-plates. Plates were then placed at 37°C for 20 min. to allow the Matrigel to polymerize before adding 500 μL of medium per well composed of 49.5% Intesticult OGM Human Basal Medium (StemCell) + 49.5% Organoid supplement (StemCell) supplemented with 10 μM Y27632 and 1% PSA. Organoids were cultured in an incubator at 37 °C, 5% CO_2_ and 90% humidity. The medium was removed after 24 h and replaced by 49.5% Intesticult OGM Human Basal Medium + 49.5% Organoid supplement + 1% PSA, which was replaced every 48h.

Organoids were passaged at a 1 : 4 ratio between the 7th and 15th day of primary culture when they presented a sufficient size (>150 µm) and some of them harbored a budding appearance. The medium culture was replaced by 750 μL of Gentle Cell Dissociation Reagent (GCDR, Stem Cell). Matrigel domes were mechanically dissociated by scratching and pipetting the GCDR solution. The operation was repeated once in order to recover all the organoids from the well. The tubes containing the GCDR solution + organoids were incubated on a shaking plate for 10 min. at 40 rpm at room temperature then centrifuged at 290 g, 4°C for 5 min. The organoids were then suspended in a solution of DMEM-F12 (Gibco) + 1% PSA + 1% fetal bovine serum (FBS), filtered at 70 μm and resuspended into a new Matrigel dome as described above. All organoids were cultured for a minimum of 3 passages.

### RNA extraction and real time RT-qPCR analysis

RNA was extracted from colonic and jejunum organoids derived from 7-day old piglets after 2 passages. Four organoid wells were harvested per condition on the 8^th^ day of culture by dissolving the Matrigel dome in a GCDR solution as described for organoid passaging. After centrifugation at 290 g for 5 min. at 4° C, organoids were rinsed several times with cold PBS until complete removal of Matrigel. Organoids were pelleted and stored at −80° C until RNA extraction using the RNeasy Micro Kit (Qiagen). Reverse-transcription was performed with 850 ng of RNA using the High Capacity cDNA RT kit (Applied Biosystems). Gene expression analysis was carried out using the Wafergen Smartchip cycler and Smartchip Multisample Nanodispenser (Biogenouest Genomics and the EcogenO core facility, Rennes, France). A house-designed dedicated set of porcine primers already described [16] was used, which allowed the analysis of the expression of 84 genes targeted on specific intestinal functions and of 12 housekeeping genes. The targeted genes were relative to the immune system, barrier function, endocrine function, digestion/nutrient transporters and tryptophan metabolism pathways. The steadiest housekeeping genes (*aldoa*, *ywhaz* and *ppia*) were used (geometric mean of Ct) to normalize data expression of targeted genes which was expressed using the 2^−ΔΔ*Ct*^ method, with jejunal organoids as reference.

### Morphometric analysis

Morphometric analysis of organoids was performed using a ZEISS Axio Vert.A1 inverted microscope equipped with an AxioCam MRC camera and a 10x objective. We took photos at day 1, 3, 6 and 8 of culture (except for the 3^rd^ subculture where cultures were stopped at day 6) of the wells in 4 constant quadrants per well and 2 heights (Fig 1B). Images were analyzed using ImageJ software. For each image, we reported the average spheroid surface (μm^2^), the average organoid (defined as structures with at least one bud) surface (μm^2^) and the percentage of organoids vs. spheroids. The results of the 16 images per condition (4 quadrants x 2 heights x 2 wells) were averaged.

### Immunohistochemistry staining

Immunohistochemistry (IHC) analysis was performed on organoids derived from 7- and 180-day old pigs after 3 passages. Matrigel domes containing the organoids were collected after 6 days of culture. The samples were fixed in a 4% formalin solution for 24 h at 4°C then in a 30% sucrose + 0.1% sodium azide solution for at least 48 h at 4°C. Matrigel domes were then coated in an OCT Embedding matrix (361603E, VWR) and stored at −20°C. 10 μm-thick sections were cut and deposited on slides stored at −20°C.

We performed immunostaining for progenitor (Sox 9), apoptotic cells (CASP3) and differentiated cells (Chromogranin A (ChgA), carbonic anhydrase (CA2), sucrase-isomaltase (SI), Villin, MUC2) identification. Slides were first thawed and rinsed for 15 min. in PBS on a shaking plate. For IHC protocols that required an unmasking step, slides were placed 15 min in a Tintoretriver-Pressure-Cooker, Low pression 110°C with an EDTA buffer pH9 1x (Diagomics BSB0030), except for ChgA where the buffer was citrate buffer pH 6 1x (Diagomics BSB0020). Slides were then rinsed with PBS for 15 min. and blocked and permeabilized with a 0.2% Triton X-100 (Merck T8787) - 3% horse serum (Merck, ref H1270) PBS solution for 1 h. The slides were rinsed with PBS then 50 μL of primary antibody (S1 Table) was deposited per section for 2 h., 50 μL of secondary antibody were deposited after PBS washing for 1.5 hours at room temperature. Finally, the sections were rinsed for 15 min. in PBS and then mounted under a coverslip using a mounting medium with DAPI (Aqueous Fluoroshield, Abcam104139). Images were obtained using a Zeiss microscope Axio Imager M2 with an ApoTome 2 module. The objective lense used was x20. Images were analyzed using ImageJ software. All immunostaining protocols were first validated on jejunal and colonic tissues (S1 Fig).

### Statistical analysis

Data are presented as means ± standard of the mean. qPCR data were analyzed by Kruskal-Wallis test using GraphPad Prism 6.07. Partial Least Square – Discriminant Analysis (PLS-DA) was performed using Metaboanalyst 5.0. Growth parameters and immunostaining data were analyzed using two-way ANOVA (age and site) after log-transformation of data. Significant differences were considered for P<0.05.

## Results

### Establishment of porcine jejunal and colonic organoids cultures

We generated jejunum and colonic organoids for all 20 pigs. For two 28-day old pigs, we did not obtain any jejunum organoids after 15 days of primary culture. For 2 pigs (aged 28 and 36 days), organoids had to be frozen before the 2^nd^ subculture because of the lockdown due to the covid-19 epidemic. When thawed, the culture of colonic organoids from the 36-day old pig failed. Finally, colonic organoid cultures were contaminated for 2 of the 7-day old pigs during the 3^rd^ passage.

### Gene expression of 3D organoids was influenced by intestinal tissue origin

We first sought to confirm that organoids derived from jejunum or colon displayed distinct phenotype at the gene expression level. We analyzed the expression of more than 80 genes in organoids derived from jejunum and colon of 7-day old piglets and passaged twice. Fifteen genes were differentially expressed between jejunal and colonic organoids (Fig 2A and S2 Table). Among the most striking differences, lyzozyme (*lyz*), Reg3g (*reg3g*), Pept-1 (*slc15a1*) and Gpr43 (*ffar2*) gene expression was greater in jejunal than in colonic organoid while carbonic anhydrase 1 (*ca1*) and toll-like receptor 2 (*tlr2*) gene expressions were merely absent in jejunal organoids compared to colonic ones (Fig 2B). PLS-DA confirmed distinct gene expression signatures between colonic and jejunal organoids (Fig 2C), that were driven by *lyz*, *slc15a1*, *ca1*, *reg3g*, *ffar2* and *slc1a1* gene expression (Variable Importance in Projection score >2, Fig 2D and S3 Table).

**Fig 2:**
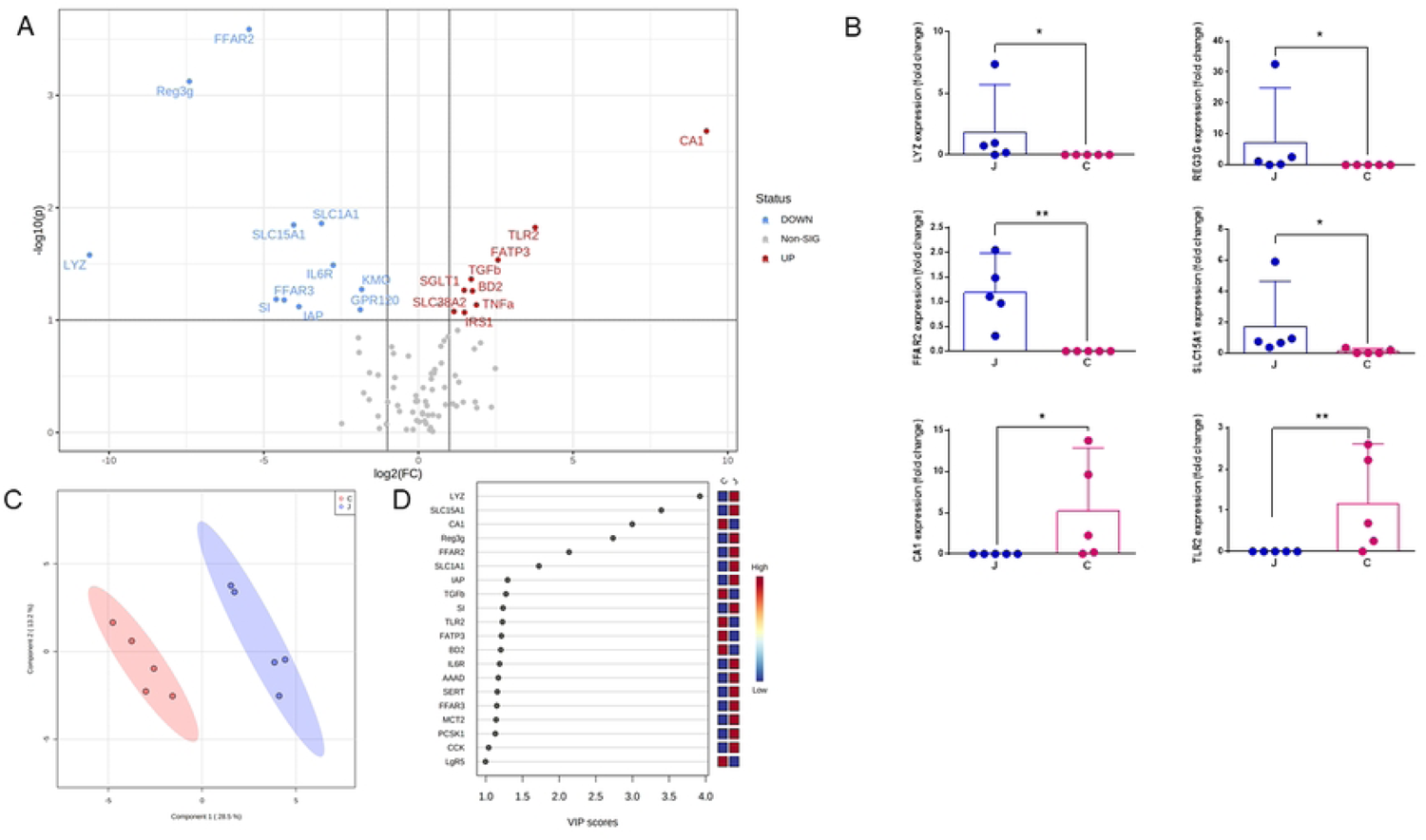
Gene expression differed between jejunal and colonic organoids. Jejunal (J) and colonic (C) organoids originating from 7-day old pigs were cultivated and passaged twice to evaluate the relative expression of 80 genes. **(A)** Volcano plot of differentially expressed genes between colonic and jejunal organoids (red: up-regulated and blue: down-regulated in colonic organoid compared to jejunal ones, FC >1.4 and P<0.05). **(B)** Relative gene expression of some specific markers of jejunal and colonic organoids. **(C)** PLS-DA and (D) VIP of relative expression of genes. * P<0.05

### Donor age influenced organoid growth

We next analyzed the effect of pig age and intestinal location on morphometric parameters of spheroids and organoids at day 6 of the primary culture and of the 1^st^ and 3^rd^ subcultures. Data presented are those at day 6 only since the 3^rd^ subculture was stopped at that time but all data are presented in S2 Fig. At day 6 of culture, the ratio between organoids and spheroids varied with pig age for the primary culture and the 3^rd^ subculture and with initial intestinal site for the primary culture and the 1^st^ subculture (Fig 3A-C). We also analyzed data by grouping data from pigs that did not had access to solid feed, i.e. 1 and 7-day old piglets, *vs* those that did have access, i.e. 28, 36 and 180-day old pigs, assuming that the luminal environment would differ [17,18]. Interestingly, the percentage of organoids to spheroids in the primary culture was greater for organoids originating from younger pigs that did not had access to solid feed than older ones that did (Fig 3D) and this ratio was maintained with increasing number of passages (Fig 3F and G). The percentage of organoids to spheroids was higher in colonic cultures than jejunal culture for the primary culture and the 1^st^ subculture (Fig 3D and E).

**Fig 3:**
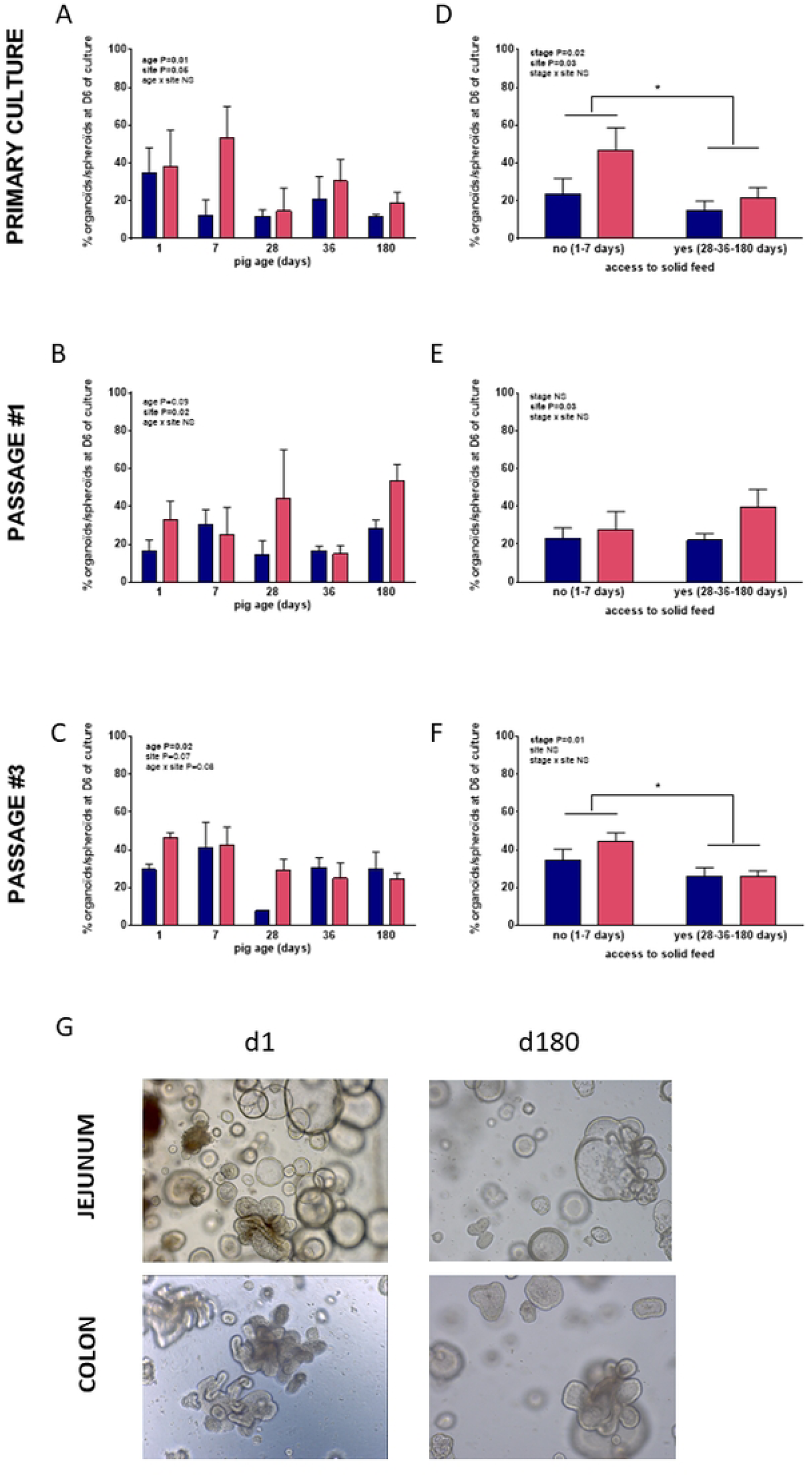
Effect of pig age on organoid to spheroid ratio at different passages. Jejunal (blue) and colonic (pink) organoids were prepared from crypts collected on pigs aged 1 to 180 days and their morphology examined using 2D-pictures after 6 days of culture during the primary culture **(A)**, first **(B)** and third **(C)** passages to evaluate the ratio of budded organoids to spheroid. Data were also stratified upon pig access to solid feed (primary culture **(D)**, first passage **(E)**, third passage **(F)**). **(G)** Representative pictures of jejunal and colonic spheroids and organoids originated from young (d1) and older (d180) pigs and passaged three times. * P<0.05

Spheroid surface varied with pig age for all cultures (tendency P=0.06 for the primary culture, Fig 4A-C). When analyzing the data by separating pigs upon their access to solid feed, we observed a stage effect that reversed with the number of passages: spheroids originating from piglets that had access to solid feed where smaller during the primary culture (P=0.002, Fig 4D and G) but bigger at passage #3 (P=0.02, Fig 4F and G). Spheroid surface was not influenced by the intestinal location, for any of the culture.

**Fig 4:**
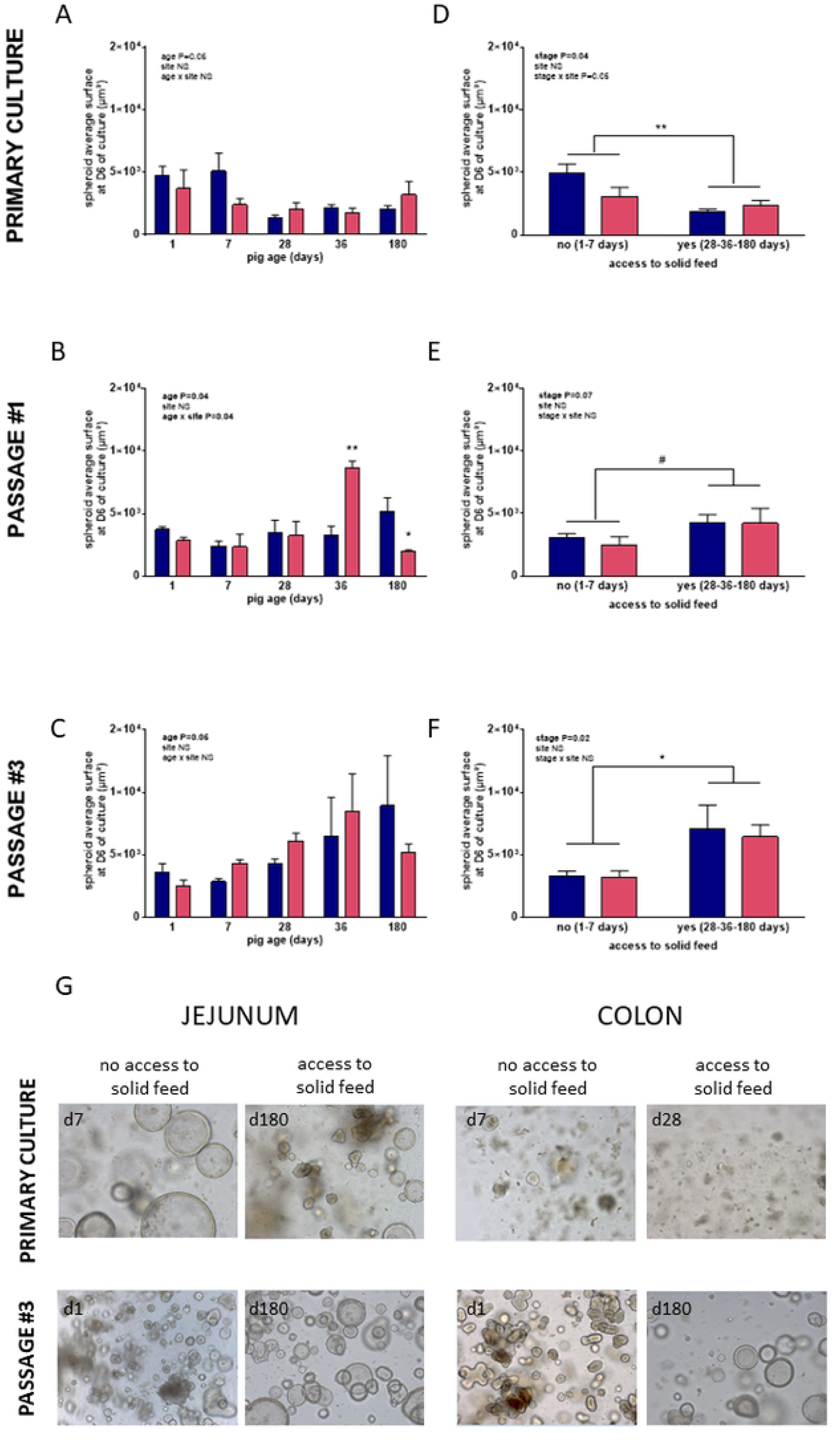
Effect of pig age on spheroid surface at different passages. Jejunal (blue) and colonic (pink) organoids were prepared from crypts collected on pigs aged 1 to 180 days and their morphology examined using 2D-pictures after 6 days of culture during the primary culture **(A)**, first **(B)** and third **(C)** passages to evaluate spheroid surface. Data were also stratified upon pig access to solid feed (primary culture **(D)**, first passage **(E)**, third passage **(F)**). **(G)** Representative pictures of jejunal and colonic spheroids originated from pigs that had access or not to solid feed, during the primary culture or the third passage. * P<0.05

Organoid surface was influenced by pig age for the 1^st^ subculture (Fig 5B and H) and by intestinal location for the primary culture and the 3^rd^ subculture (Fig 5A, C and G). These effects were also observed when grouping piglets upon their access to solid feed: during the primary culture, organoids from young pigs were larger than that from older pigs (Fig 5D). This effect was reversed for the 1^st^ subculture (Fig 5E). For the primary culture and the 3^rd^ subculture, colonic organoids were bigger than jejunal ones (Fig 5E).

**Fig 5:**
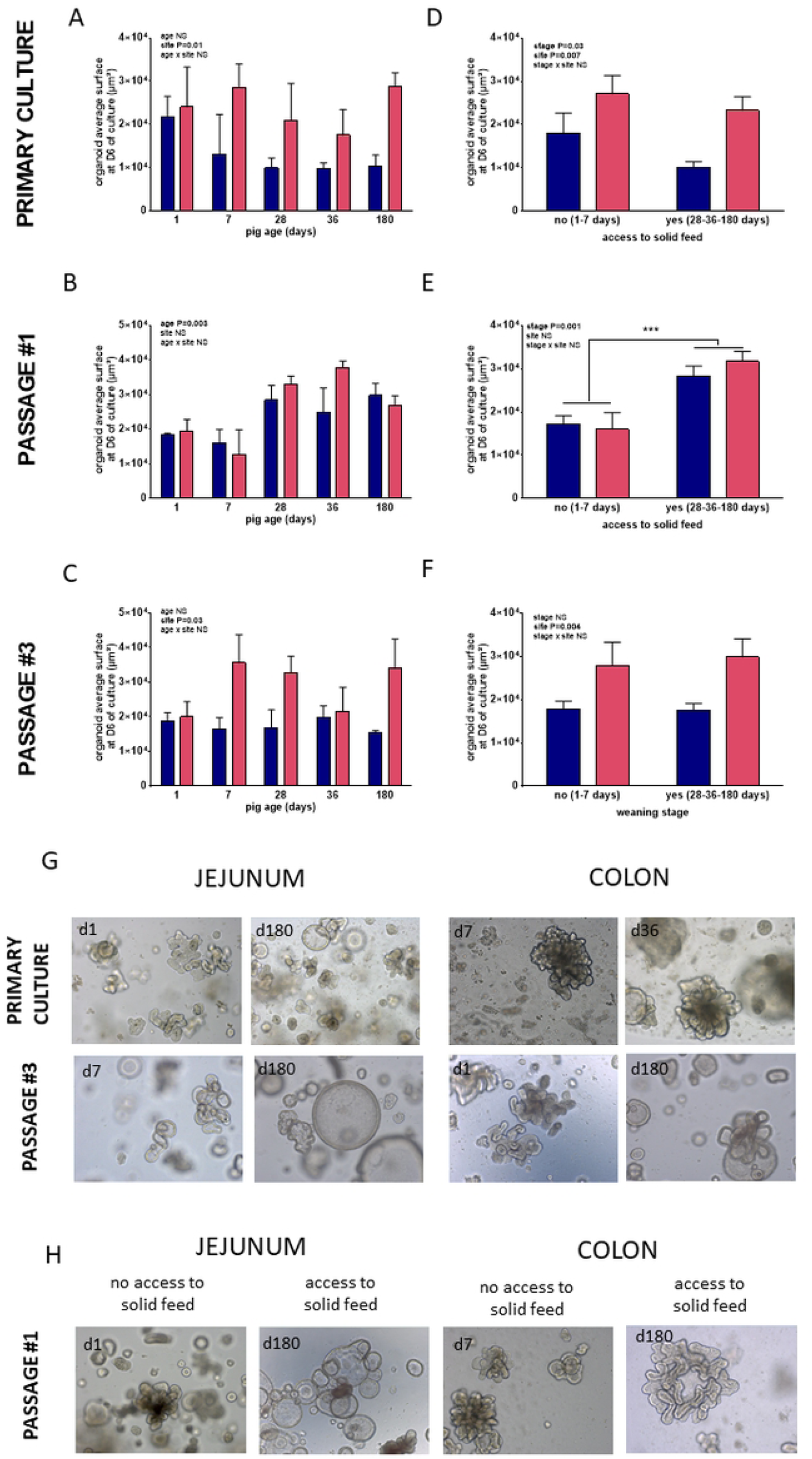
Effect of pig age on organoid surface at different passages. Jejunal (blue) and colonic (pink) organoids were prepared from crypts collected on pigs aged 1 to 180 days and their morphology examined using 2D-culture after 6 days of culture during the primary culture **(A)**, first **(B)** and third **(C)** passages to evaluate organoid surface. Data were also stratified upon pig access to solid feed (primary culture **(D)**, first passage **(E)**, third passage **(F)**). **(G)** Representative pictures of jejunal and colonic organoids during the primary culture or the third passage. **(H)** Representative pictures of jejunal and colonic organoids, originating from that had access or not to solid feed, during the second passage. *** P<0.001

### Donor age did not influence organoid cellular composition

In order to evaluate whether pig age had an effect on organoid cellular composition, we performed immunostaining analysis of jejunal and colonic organoids from 7 and 180-day old pigs. Villin immunostaining was evident in all organoids with an apical-in polarity, irrespective of the pig age and intestinal location (Fig 6A).

**Fig 6:**
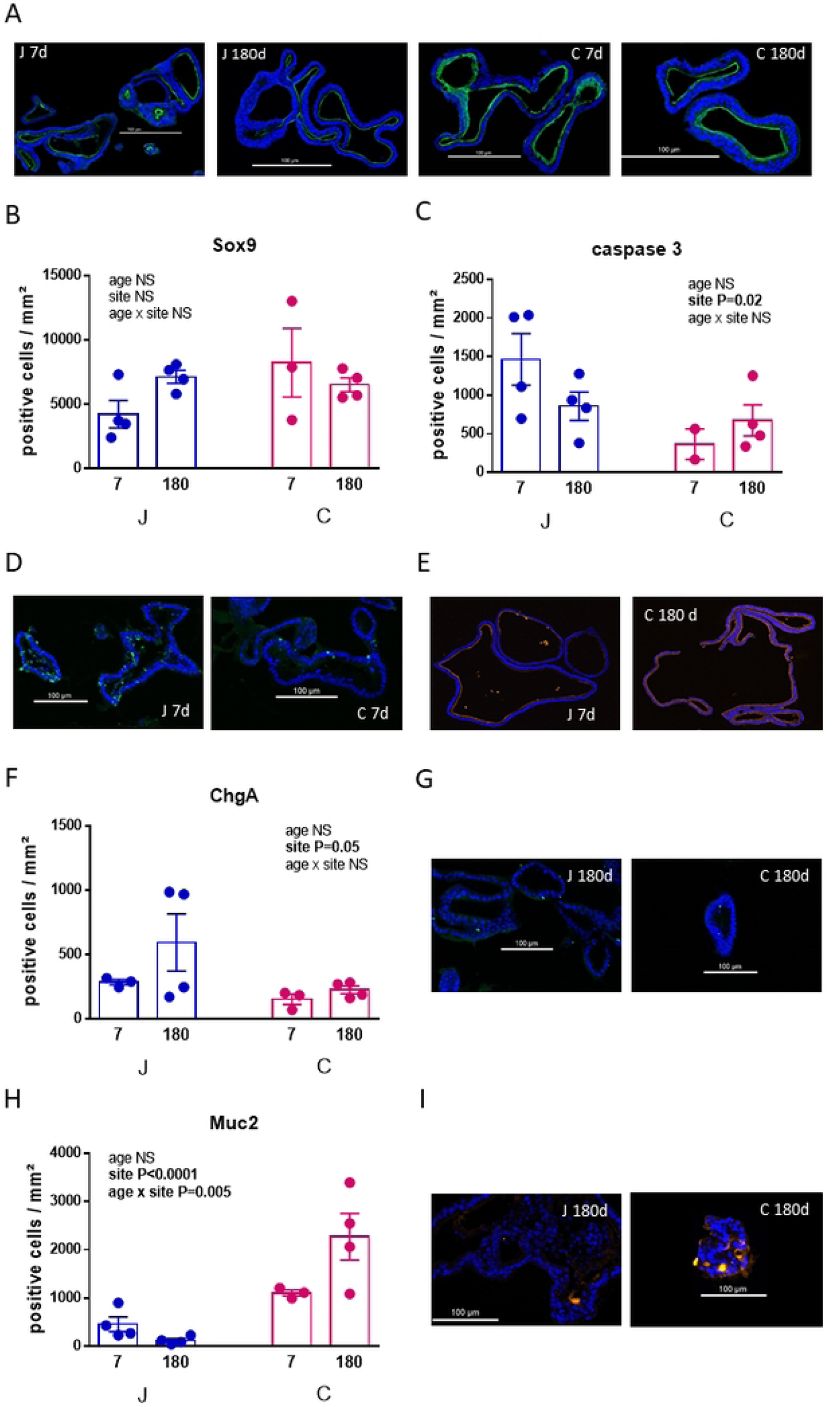
Effect of pig age on organoid cellular composition. Jejunal (blue) and colonic (pink) organoids were prepared from crypts collected on pigs aged 7 or 180 days and immuno-staining for villin (**A:** representative pictures), Sox9 **(B)**, caspase 3 (**C, D:** representative pictures), CA2 (**E,** representative pictures), chromogranin A (**F, G**: representative pictures) and Muc2 (**H, I:** representative pictures) was performed on fixed organoids after 3 passages. * P<0.05

The average density of Sox9-positive cells did not differ between organoids originating from 7 or 180-day old pigs, irrespective of the intestinal location (Fig 6B). On the contrary, the density of caspase 3-positive cells was lower in organoids originating from colon than jejunum, irrespective of the pig age (Fig 6C and D).

Then we evaluated gene expression of specific protein representing IEC types: carbonic anhydrase II (colonocytes), sucrase isomalatase (enterocytes), chromograninA (entero-endocrine cells) and mucin 2 (goblet cells). The CA II protein was expressed at the apical side of colonic organoids but surprisingly also of jejunal ones (Fig 6E). This immunofluorescent labeling was sometimes heterogeneous, homogeneous or absent. We did not highlight any difference according to the age of pigs or tissue (data not shown). We did not observe any immunofluorescent labeling for the SI protein (data not shown). However, tests previously performed on colonic and jejunal tissues clearly showed presence of immunofluorescence in jejunal tissue and absence in colonic ones (S1 Fig). Chromogranin-A was used to label entero-endocrine cells. The density of ChgA-positive cells was higher in jejunal organoids than colonic ones, irrespective of pig age (Fig 6F and G). Finally, Muc2-positive cell density was significantly higher in organoids originating from colon than jejunum. The density of Muc2-positive cells decreased with age in jejunal but not colonic organoids (Fig 6H and I).

## Discussion

Getting a better understanding of the impact of culture methods and ISC donor characteristics on intestinal organoid is essential to encourage the use of this alternative to animal experiment. Our objective was therefore to evaluate if the age of the ISC donor influences organoid growth, morphology and phenotype in the porcine species. We established that pig age influenced organoid growth and budding (*i.e.* proliferative and morphologic processes) with enhanced growth and budding when ISC were collected in young animals compared to older ones. Hence, cellular composition (*i.e*. differentiation processes) was barely affected by ISC donor age. Noteworthy, some of these age effects on organoid growth differed between primary cultures and passaged organoids, indicating that the culture medium modified this developmental programming.

The neonatal period is known as an intensive growth period of the intestine in pig, along with body weight increase [19]. This massive growth is due to intense continuous self-renewal of ISC followed by daughter cell transit and differentiation along the crypt villus-axis. Milk-derived nutrients and growth factors are believed to be strong drivers of ISC proliferation during this period [20]. Microbiota implantation also plays a role, likely reducing ISC proliferation, since germ-free piglets harbors longer villi compared to conventional ones, especially in the distal small intestine [21]. Here, we observed heightened growth of organoids derived from young compared to older pigs, despite similar culture conditions, *i.e.* similar growth factor concentrations and absence of bacterial stimuli. Even if we do not know the exact composition of the commercial medium, this later likely contains high concentration of proliferative factors as it is meant to keep organoid in a proliferation state. This could suggest that ISC sensitivity to growth factors is reduced when ISC are derived from old pigs compared to young ones. Our protocol of organoid generation is based on crypt but not isolated ISC seeding in Matrigel. Hence, a different number of ISC per crypts could account for difference in organoid growth. However, both a reduced number and an increased number of ISCs has been described in aged mice compared to young ones, rending difficult to conclude on the effect if age on ISC number. Yet, all studies agree on reduced ISC function upon aging in mice, due to reduced canonical Wnt signaling and altered Notch signaling [22]. To our knowledge, no data on pig ISC senescence are available. Moreover, adult pigs we used in our study were young adult pigs aged 180 days, in which aging processes described above are probably not fully committed yet. Thus, further studies are needed to fully understand the effect of age on ISC function in pigs.

Another factor that was influenced by ISC donor age in our study was the budding capacity of organoids. Indeed, the ratio of budded organoids to spheroids was greater when ISC were derived from young compared to old pigs. It has been suggested that morphogenesis of budding intestinal organoids results from either locally enhanced cell proliferation [23], or biomechanical processes like growth-induced buckling [24] or from internal pressure combined with higher cell contractility in the regions containing differentiated cells compared to the regions containing ISC [25]. Budding of organoids is therefore not a real marker of differentiation but rather of enhanced proliferation or onset of differentiation. Thus, the difference observed in the budded organoid to spheroid ratio between young and older pig - derived organoids in our study is likely supported by difference in proliferative rates of ISC as discussed above. Another result of our study is the fact that while spheroids originating from young piglets were larger than those originating from older pigs during primary culture, this effect reversed with organoid passaging. This plasticity of ISC under identical culture conditions suggests changes in ISC programming, a phenomenon likely due to epigenetic drift [26], which might differ upon initial ISC donor age. Understanding the mechanisms underlying this drift was out of the scope of our study but such growth potential changes with organoid passaging should be acknowledged when cultivating organoids.

Interestingly, one factor that seemed to reduce organoid growth was the fact that piglets were submitted or not to solid feed. Indeed, while 1- and 7-day old piglets were only fed sow milk, 28-day old piglets probably ate a mix of sow milk and creep-feed, that was available in their pen from day 21 of life. Creep feed intake is variable among piglets and is driven by both homeostatic needs and exploratory behavior. Indeed, milk production by the sow starts to plateau after 3 weeks of lactation, forcing piglets to find alternative nutrient supply [27]. Moreover, exploratory behavior, *e.g*. nosing, rooting, chewing, and object play peaks around four weeks of age [28]. Thus, although we did not measure individual solid feed intake, we are confident that all piglets ate some of the creep feed available in their pens. Introduction of solid feed induces tremendous change in gut luminal environment. In rabbits, where the unique feeding behavior of the species allows to experimentally disentangle the effect of solid feed introduction from that of developmental processes, it has been shown that the caecal microbiota composition and metabolome of rabbits fed both milk and solid feed was similar to that of age-matched rabbits fed a solid diet only and largely differed from that of age-matched milk-fed rabbits [17]. Similarly, in pigs, introduction of solid feed dramatically changes the luminal microbiota and metabolome [29]. As mentioned earlier, the microbiota interferes with epithelial cell proliferation and differentiation [30]. Bacterial metabolites, including short-chain fatty acids and bacterial components have been shown to modulate ISC activity. Butyrate and propionate reduce ISC proliferation in organoids [31,32], while LPS enhanced dog intestinal organoid proliferation [33]. *In vivo*, luminal concentration of short chain fatty acids, including butyrate and propionate increases with piglet age, while that of LPS probably decrease due to the reduced relative abundance of Proteobacteria with age [13]. Hence, we speculate that these changes at the luminal level programs ISC that harbor a less proliferative capacity when cultivated *in vitro*.

Cellular composition of organoids derived from young versus old pigs did not differ in our study, suggesting no impact of pig age on the differentiation process. We chose not to induce differentiation by changing medium composition in our study, since our objective was to passage organoids several times. Yet, using IHC techniques to specifically target proteins, we observed the presence of several proteins specific of differentiated cells, such as chromogranin A and Muc2, markers of secretory cell lineage or CA2, specific of colonocytes. Noteworthy, while *ca2* gene expression was restricted to colonic organoid and IHC revealed no CA2 staining in pig jejunal sections, CA2 protein was observed in organoids derived from both jejunal and colonic ISC. Similarly, whereas *SI* gene expression was restricted to jejunal organoid and SI staining was observed only in jejunal, but not colonic sections, no SI staining was observed in organoids, irrespective of their intestinal origin. To date, we have no explanation for such inconsistencies. Yet, despite these discrepancies, our IHC data revealed that spontaneous differentiation seemed to have occurred in our culture conditions. *In vivo*, the differentiation process of transit amplifying cells is orchestrated by the dynamic equilibrium of several pathways, including Wnt, bone morphogenic protein, Notch and mitogen-activated protein kinase pathways, which form distinct gradients along the crypt–villus axis. This equilibrium between growth and differentiation factors ultimately guides the cell fate specification toward mature cell types. In organoid cultures, such gradients do not exist. We speculate that proliferation and growth process of organoids move away daughter cells from the stem niche, reducing the influence of niche factors, likely allowing some differentiation. Protocols involving reduction of the growth factor concentration in the culture medium induce more profound differentiation processes, with reduced Lgr5 expression and enhanced expression of differentiated cell markers [4]. Hence, further studies using such protocols are needed to fully conclude on the absence of ISC donor age on epithelial cell differentiation.

In conclusion, our aim was to provide data on porcine intestinal organoid proliferation and differentiation processes upon initial ISC donor age, to encourage the use of this *in vitro* model in further studies. Our study shows the importance of taking into account the donor age and passage number when intestinal organoids are used to evaluate intestinal epithelial cell proliferation processes. On the other hand, if the objective is to work on differentiation processes, the pig age does not seem to be at play.

## Author contribution

CD, GR, LLN, VR, SL: investigations, CD, APA and GB: conceptualization, funding acquisition, formal analysis, original draft preparation, all authors; review and editing

## Supporting information

**S1 Table: List of antibodies used for immuno-histochemistry protocols**

**S2 Table: Relative gene expression of organoids derived from jejunum or colon after two passages**

**S3 Table: Variable Importance in Projection Score for each variable of the PLS-DA**

**S1 Fig: Representative pictures of IHC staining on jejunal and colonic tissues**

**S2 Fig: Evolution of the organoid / spheroid ratio, spheroid surface and organoid surface with days of culture**

## Notes

### Competing Interest Statement

The authors have declared no competing interest.

